# Aberrant splicing exonizes *C9ORF72* repeat expansion in ALS/FTD

**DOI:** 10.1101/2023.11.13.566896

**Authors:** Suzhou Yang, Denethi Wijegunawardana, Udit Sheth, Austin M. Veire, Juliana M. S. Salgado, Manasi Agrawal, Jeffrey Zhou, João D. Pereira, Tania F. Gendron, Junjie U. Guo

## Abstract

A nucleotide repeat expansion (NRE) in the first annotated intron of the *C9ORF72* gene is the most common genetic cause of amyotrophic lateral sclerosis (ALS) and frontotemporal dementia (FTD). While C9 NRE-containing RNAs can be translated into several toxic dipeptide repeat proteins, how an intronic NRE can assess the translation machinery in the cytoplasm remains unclear. By capturing and sequencing NRE-containing RNAs from patient-derived cells, we found that C9 NRE was exonized by the usage of downstream 5ʹ splice sites and exported from the nucleus in a variety of spliced mRNA isoforms. *C9ORF72* aberrant splicing was substantially elevated in both C9 NRE^+^ motor neurons and human brain tissues. Furthermore, NREs above the pathological threshold were sufficient to activate cryptic splice sites in reporter mRNAs. In summary, our results revealed a crucial and potentially widespread role of repeat-induced aberrant splicing in the biogenesis, localization, and translation of NRE-containing RNAs.

## INTRODUCTION

Instability and expansion of short tandem repeat (STR, also known as microsatellite) sequences in the human genome cause more than forty genetic disorders^1–3^, many of which affect the nervous system. While some nucleotide repeat expansions (NREs) such as (CAG)_n_ repeats in Huntington’s disease reside within protein coding sequences and cause disease primarily by producing repetitive polypeptides such as poly-glutamine^4^ (polyQ), the pathogenic mechanisms of NREs within noncoding regions, including 5ʹ untranslated regions (UTRs), introns, and 3ʹ UTRs, are much less well understood. Some noncoding NREs, such as (CTG)_n_ and (CCTG)_n_ repeats in myotonic dystrophy type 1 and 2 (DM1/2), respectively, are thought to cause cellular dysfunction by producing toxic repeat RNAs that can sequester RNA-binding proteins^5,6^. In addition, several “noncoding” NREs have been shown to produce repetitive polypeptides despite the lack of an in-frame upstream AUG codon via a poorly understood mechanism now known as repeat-associated non-AUG (RAN) translation^7^. Initially discovered in myotonic dystrophy type 1 (DM1) and spinal cerebellar ataxia type 8 (SCA8)^ref.8^, which are caused by (CTG)_n_ NREs in *DMPK* and *ATXN8* genes, respectively, the list of NREs that undergo RAN translation now includes those within annotated protein coding sequences (e.g., Huntington’s disease^9^), 5ʹ UTRs (e.g., fragile X-associated ataxia syndrome^10^), and perhaps most surprisingly, introns (e.g., DM2^ref.11^ and C9 ALS/FTD^12–14^). When ectopically expressed in cellular and animal models, many RAN-translated polypeptides can recapitulate various pathological features and cause cellular dysfunction^7,15–22^.

The most common genetic cause of ALS and FTD, accounting for nearly half of the familial cases and ∼4% of sporadic cases, is a (GGGGCC)_n_ NRE within the first annotated intron of the *C9ORF72* gene^23,24^. Similar with other disease-associated NREs, multiple pathophysiological mechanisms have been proposed for C9 ALS/FTD^25,26^. The C9 NRE is transcribed in both sense and antisense directions, producing at least two types of NRE RNAs, each of which can form nuclear foci and recruit a variety of RNA-binding proteins^12,14,24,27^. In addition, both sense and antisense C9 NRE RNAs undergo RAN translation^12–14^, with all six types of dipeptide repeat (DPR) proteins having been detected in patient tissues at varying abundance. Among them, the arginine-rich DPRs, including the poly(glycine-arginine) (polyGR) encoded by the sense NRE and poly(proline-arginine) (polyPR) by the antisense NRE, can cause acute toxicity when ectopically expressed in cells in part by globally inhibiting translation^19,21,28^, translation-dependent mRNA surveillance^21,29,30^, ribosome biogenesis^15,20^, and nucleocytoplasmic export^15,16,31^.

Despite the accumulating evidence supporting a pathophysiological role of C9ORF72 RAN translation, the question remains how an intronic NRE can gain access to the translation machinery. The initial model has been that the NRE may inhibit splicing and cause the retention of *C9ORF72* intron 1^ref.32–34^, which is more than 6kb in length (not including the NRE) and contains multiple putative polyadenylation signals. More recently, a study using SunTag-based translation reporter systems suggests a drastically different model in which the excised intron 1 lariat is stabilized by C9 NRE, exported from the nucleus, and translated in a cap-independent manner^35^. Partly due to the low abundance of NRE-containing RNA in question, direct characterization of these RNAs has proven challenging. To overcome this challenge, we devised an unbiased method to enrich low-abundance NRE-containing RNAs from patient-derived cells by using antisense oligonucleotides (ASOs) and performed high-throughput RNA sequencing to delineate their sequence composition. Our results revealed a distinct “exonization” mechanism in which C9 NRE was retained as part of an extended exon in a variety of splice isoforms. This mechanism provides a parsimonious explanation for the retention of intronic NREs in a readily exported and translatable mRNA and has broad implications for other disease-associated NREs.

## RESULTS

### ASO-based capture of NRE-containing transcripts

Conventional RNA-seq analysis cannot distinguish reads from the NRE-containing transcripts from the much more abundant C9ORF72 mRNAs without NRE. To specifically enrich the NRE-containing transcripts, we adapted the previously described RNA antisense purification (RAP) method^36^ and used 5ʹ-biotinylated (CCCCGG)_3_ ASOs to capture (GGGGCC)_n_ NRE-containing transcripts from cytoplasmic RNAs of patient-derived fibroblasts for RNA-seq analysis (hereafter referred to as NRE-RAP-seq, **Fig. 1a**). In addition to two C9 NRE^+^ fibroblast lines (FB504 and FB506), two C9 NRE^−^ fibroblast lines were used as negative controls, including FB485 with a *MAPT* P301L mutation and FB503 with no identified genetic risk factors. After hybridization, the RNA:DNA duplexes were pulled down using streptavidin-conjugated magnetic beads. After stringent washing, the NRE-containing transcripts were eluted by RNase H digestion and used for RNA-seq.

**Figure 1.**
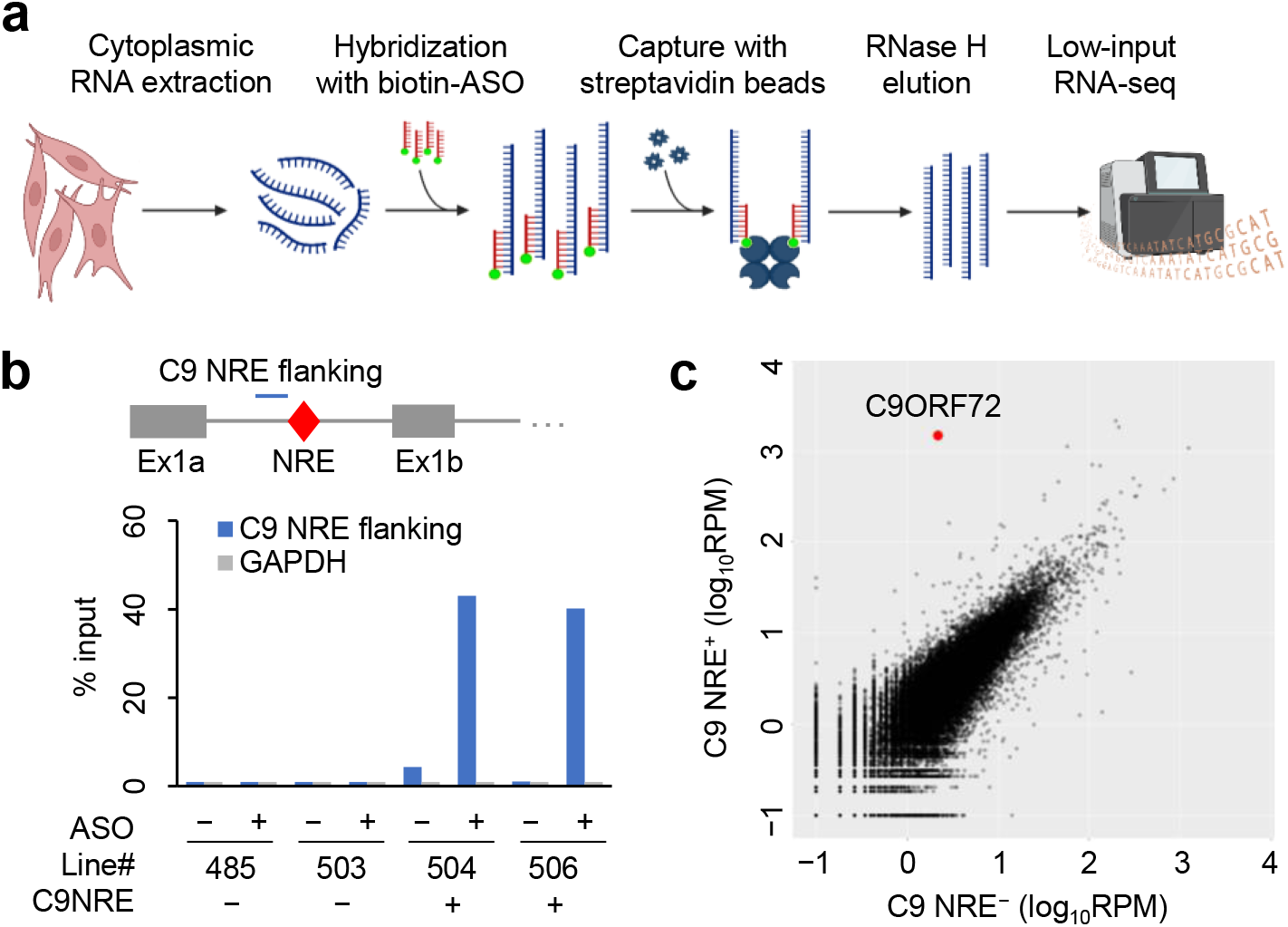
ASO-based capture of NRE-containing transcripts. (**a**) Schematics of NRE-RAP-seq. (**b**) RT-qPCR quantification of the enrichment of C9 NRE-flanking region and GAPDH by NRE-RAP relative to input. (**c**) Comparison of normalized NRE-RAP-seq read counts for each gene between C9 NRE^+^ and NRE^−^ samples. C9ORF72 is indicated in red.

To confirm the specific enrichment of C9 NRE-containing transcripts by NRE-RAP, we measured the abundance of the C9 NRE upstream flanking sequence (the NRE itself cannot be quantitatively amplified by PCR) by RT-qPCR. Consistent with the notion that C9 intron 1 is fully spliced out and therefore absent in the cytoplasm of C9 NRE^−^ cells, the C9 NRE flanking region was undetectable in ASO-captured RNAs (**Fig. 1b**). In contrast, this region was highly enriched after NRE-RAP in both C9 NRE^+^ fibroblast lines (**Fig. 1b**). The enrichment strictly required the addition of (CCCCGG)_3_ ASO and was not observed for GAPDH mRNA (**Fig. 1b**).

For further validation, we quantified and compared NRE-RAP-seq read counts for each gene between C9 NRE^+^ and C9 NRE^−^ samples. The results were highly correlated between biological replicates (**Supplementary Fig. 1**). Importantly, C9ORF72 was the single most enriched transcript, yielding ∼1,000-fold more reads in C9 NRE^+^ cells than NRE^−^ controls (**Fig. 1c**), indicating high specificity and sensitivity of NRE-RAP-seq.

### Exonization of C9 NRE by downstream 5ʹ splice site usage

Close examination of the RNA-seq read coverage at the *C9ORF72* locus uncovered several critical features of cytoplasmic NRE-containing transcripts. First, similar to the cytoplasmic RNA input, a majority of reads mapped to *C9ORF72* exons (**Fig. 2a**), arguing against excised intron 1 being the predominant form of NRE-containing transcripts. Second, the distribution of reads mapping to *C9ORF72* intron 1 was inconsistent with the full retention of intron 1, with the vast majority of reads mapping to the 5ʹ half of the intron (**Fig. 2a, b**). Similar results were observed in an independent C9 NRE^+^ fibroblast line (**Supplementary Fig. 2a**). Third, analysis of reads that spanned exon-exon junctions indicated that the observed read coverage in intron 1 could be readily explained by splicing at one of many 5ʹ splice sites downstream from the NRE (**Fig. 2b**). The three most abundant splice junctions all used the canonical 3ʹ splice sites of intron 1 (**Fig. 2b**, red), with the most abundant junction using the canonical 5ʹ splice site at the end of exon 1b and the next two most abundant junctions each using a cryptic 5ʹ splice sites representing various extended forms of exon 1, which we termed exon 1c and 1d (**Fig. 2b**, red). Aside from these most abundant junctions, numerous additional unannotated splice junctions were detected in ASO-captured NRE-containing transcripts (**Fig. 2b**, grey), including some using two cryptic splice sites.

**Figure 2.**
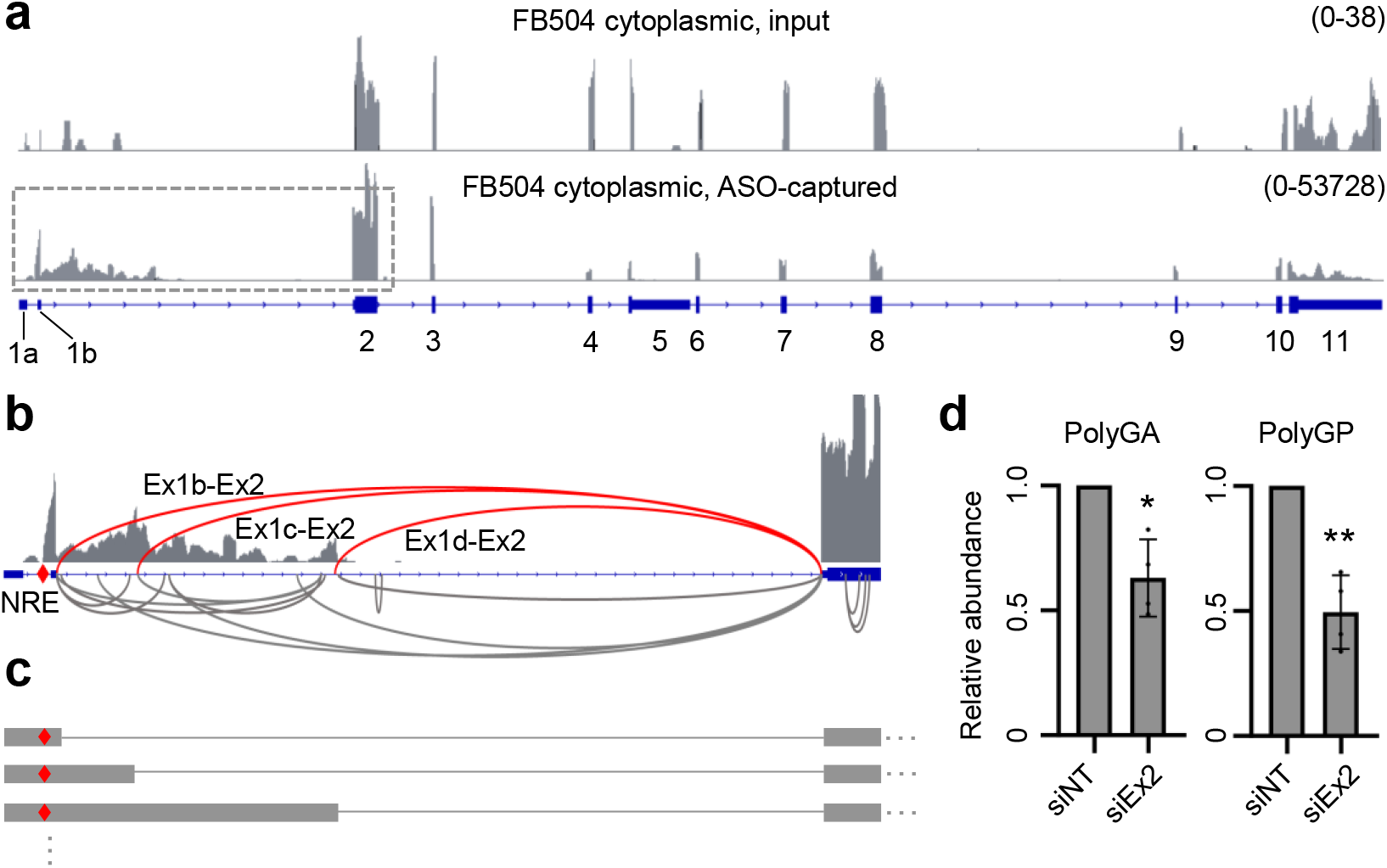
Exonization of C9 NRE by downstream 5ʹ splice site usage. (**a**) Read coverage of FB504 cytoplasmic RNA input (*top*) and ASO-captured NRE-containing RNAs (*bottom*). Dotted line rectangle indicates the region shown in (**b**). (**b**) Splice junctions within and near intron 1 detected in NRE-RAP-seq results. The three most abundant splice junctions are shown in red. (**c**) Schematics illustrating C9 NRE-exonized transcript isoforms. (**d**) Quantification of polyGA and polyGP abundance in siRNA-treated C9 NRE^+^ fibroblasts. siNT, non-targeting siRNA. siEx2, a mix of two siRNAs targeting exon 2. *, p<0.05; **, p<0.01, two-tailed ratio t test.

To further validate the connectivity between NRE and these aberrant splice junctions, we performed RT-qPCR with ASO-captured RNAs and junction-specific primers. Consistent with the RNA-seq results, the Ex1c−Ex2 splice junction was greatly enriched by NRE-RAP in an ASO-dependent and NRE-dependent manner (**Supplementary Fig. 2b**). Taken together, our results suggest that in the cytoplasm, C9 NRE is predominantly “exonized” and exists in a variety of aberrantly spliced transcript isoforms (**Fig. 2c**).

Based on NRE-RAP-seq results, a majority of cytoplasmic NRE-containing transcripts shared the usage of the canonical 3ʹ splice site of intron 1 (**Fig. 2b**). To test whether these splice isoforms were actively translated, we transfected C9 NRE^+^ fibroblasts with small interfering RNAs (siRNAs) targeting exon 2 (**Supplementary Fig. 2c**). Compared to a non-targeting control siRNA, exon 2-targeting siRNAs significantly decreased the abundance of poly(glycine-alanine) (polyGA) and poly(glycine-proline) (polyGP) DPRs (**Fig. 2d**), both encoded by the sense (GGGGCC)_n_ repeats. Notably, an independent study has reported similar results by targeting exon 2 with an artificial microRNA^37^. Together, these results are consistent with the notion that these aberrantly spliced mRNAs served as the template for DPR translation.

### Nucleocytoplasmic distribution of exonized C9 NRE

To assess the nucleocytoplasmic export efficiency of C9 NRE-exonized transcript isoforms, we measured by RT-qPCR the distribution of each NRE-containing transcript isoform between nuclear and cytoplasmic ASO-captured RNAs. The NRE-upstream flanking region and the 3ʹ half of intron 1 were predominantly nuclear, consistent with these regions largely coming from unspliced pre-mRNAs (**Fig. 3a**). In contrast, all three splice isoforms (Ex1b/1c/1d−Ex2) showed similar cytoplasmic fractions (40-60%) as the canonical C9ORF72 mRNA (**Fig. 3a**), suggesting that these C9 NRE-exonized RNA isoforms are efficiently exported.

**Figure 3.**
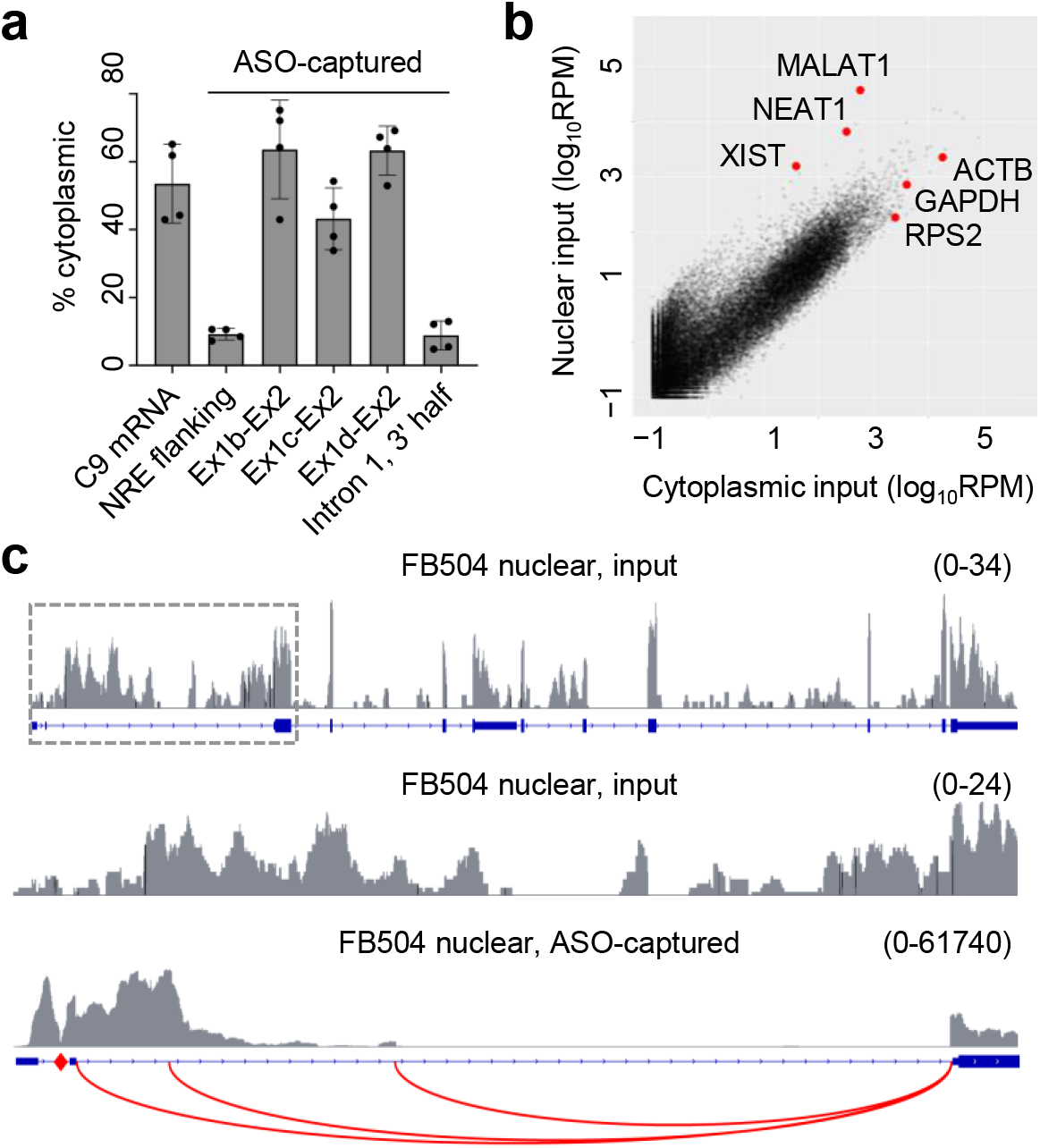
Nucleocytoplasmic distribution of exonized C9 NRE. (**a**) Comparison of normalized read count per gene between nuclear and cytoplasmic input RNAs, showing the most enriched transcripts. (**b**) Read coverage of FB504 nuclear RNA input (*top, middle*) and ASO-captured C9 NRE-containing RNAs (bottom). Dotted line rectangle indicates the region shown in the middle and bottom plots. The three most abundant splice junctions detected in NRE-RAP-seq results are shown in red. (**c**) RT-qPCR quantification of the distribution of individual C9ORF72 region between nucleus and cytoplasm. Ex2-Ex3 junction, a proxy for canonical C9ORF72 mRNA, was measured using the input RNA, whereas all other regions were measured using ASO-captured RNA.

Previous studies have shown that pre-mRNA splicing promotes nucleocytoplasmic export, whereas unspliced mRNAs are often retained in the nucleus^38^. We thus wondered whether the prevalence of spliced NRE transcripts in the cytoplasm may be in part due to inefficient nucleocytoplasmic export of other forms of NRE-containing transcripts such as unspliced mRNAs and excised introns. To test this possibility, we performed NRE-RAP-seq using the nuclear RNA fraction from C9 NRE^+^ fibroblasts. As expected, abundant nuclear noncoding RNAs (XIST, NEAT1, and MALAT1) were highly enriched in the nuclear versus cytoplasmic RNA fraction, whereas housekeeping mRNAs such as ACTB, GAPDH, and RPS2, were predominantly cytoplasmic (**Fig. 3b**).

In contrast to the cytoplasmic RNA input, which yielded predominantly exon-mapping reads, the nuclear RNA fraction yielded many more reads mapping to all *C9ORF72* introns (**Fig. 3c**), consistent with the presence of incompletely spliced pre-mRNAs. However, NRE-RAP-seq read coverage of ASO-captured NRE-containing transcripts from the nuclear fraction was similar to those from the cytoplasmic fraction in that reads mostly mapped to the *C9ORF72* exons and the 5ʹ half of intron 1 (**Fig. 3c**, **Supplementary Fig. 3**). Furthermore, Ex1b-Ex2, Ex1c-Ex2, and Ex1d-Ex2 splice junctions were also identified as the most abundant splice junctions in nuclear NRE-containing transcripts (**Fig. 3c**). These results suggest that aberrantly spliced RNAs are also the predominant form of NRE-containing transcripts in the nucleus.

### *C9ORF72* aberrant splicing in motor neurons and ALS/FTD brains

Having identified a role of aberrant splicing in the biogenesis of C9 NRE-containing transcripts in fibroblasts, we proceeded to confirm these observations in more disease-relevant contexts. We first performed RT-qPCR using total RNAs from induced pluripotent stem cells (iPSCs) and iPSC-derived motor neurons (iPS-MNs). While overall *C9ORF72* expression levels were similar between C9 NRE^+^ and NRE^−^ iPS-MNs, NRE-flanking intron RNA abundance was substantially higher in C9 NRE^+^ iPS-MNs (**Fig. 4a**). Concomitantly, aberrant Ex1c−Ex2 splice junction was increased by >20 folds in C9 NRE^+^ compared to NRE^−^ iPS-MNs (**Fig. 4a**). Similar results were also seen in undifferentiated iPSCs (**Supplementary Fig. 4**), even though overall C9ORF72 expression level was ∼10-fold lower than that in iPS-MNs.

**Figure 4.**
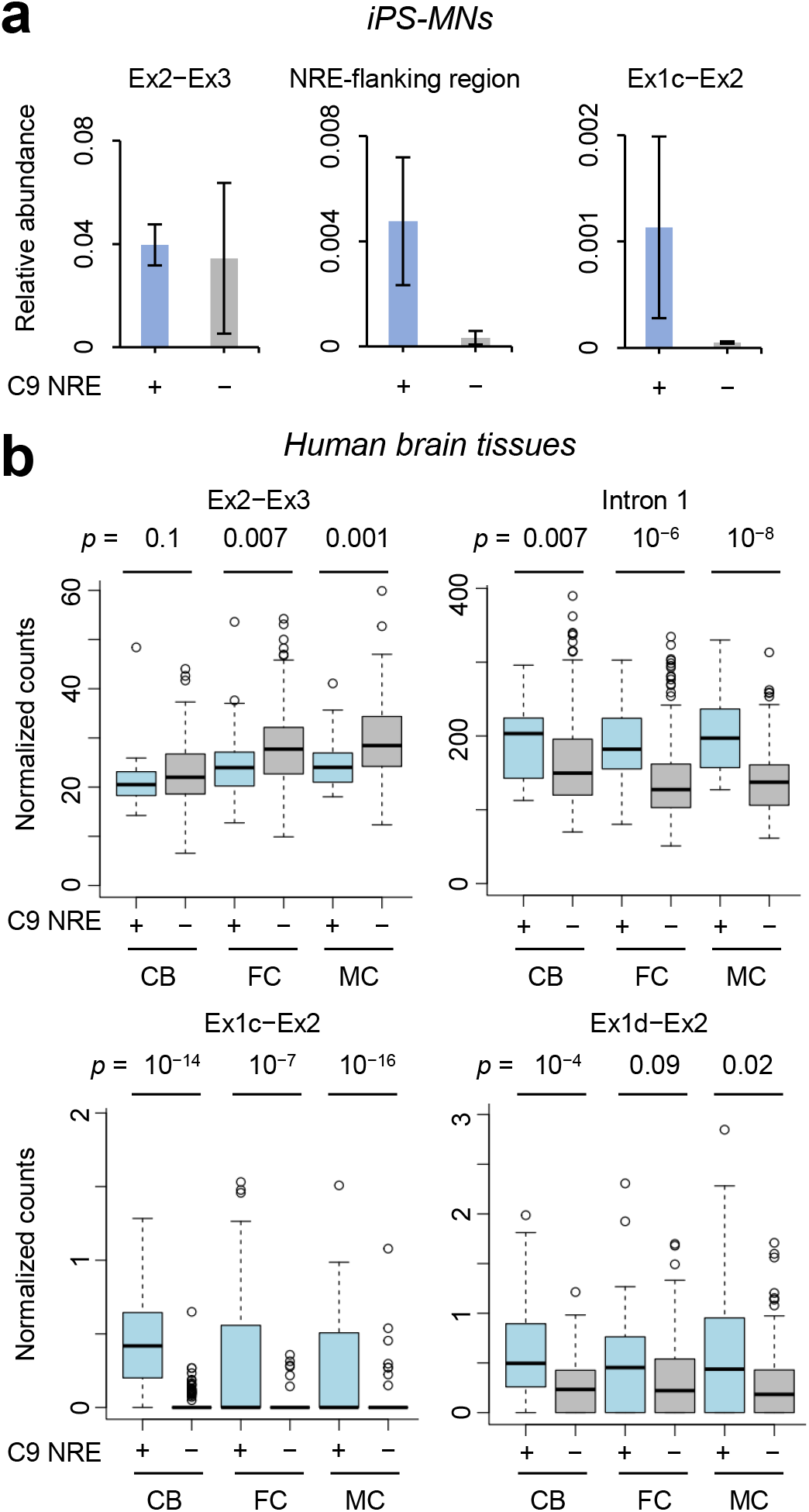
C9ORF72 aberrant splicing in motor neurons and ALS/FTD brains. (**a**) RT-qPCR quantification of the Ex2-Ex3 splice junction, NRE-flanking region, and Ex1c-Ex2 splice junction, comparing C9 NRE^+^ and NRE^−^ iPS-MNs. (**b**) RNA-seq quantification of the Ex2-Ex3 splice junction, intron 1, Ex1c-Ex2, and Ex1d-Ex2 splice junctions, comparing C9 NRE^+^ and NRE^−^ samples in each brain region. CB, cerebellum. FC, frontal cortex. MC, motor cortex. p values, Mann-Whitney-Wilcoxon tests.

We next quantified *C9ORF72* aberrant splicing events using a large RNA-seq dataset obtained through the New York Genome Center ALS Consortium, which includes 183 C9 NRE^+^ and 928 NRE^−^ tissue samples from various brain regions. In contrast to the decreased canonical *C9ORF72* Ex2−Ex3 splice junction, reads mapping to C9 NRE-containing intron 1 was significantly increased in NRE^+^ compared with NRE^−^ samples across brain regions (**Fig. 4b**). Consistent with our iPS-MN results, the aberrant Ex1c−Ex2 and Ex1d−Ex2 junctions were also significantly increased in C9 NRE^+^ cerebellum, frontal cortex, and motor cortex samples (**Fig. 4b**). Notably, Ex1c−Ex2 and the more abundant Ex1d−Ex2 junction were both detectable in some C9 NRE^−^ samples (**Fig. 4b**), albeit at lower levels compared to C9 NRE^+^ cases, suggesting that aberrant splicing can occurs spontaneously but is elevated by C9 NRE.

Since the nuclear clearing of TDP-43, a known suppressor of cryptic splice site usage, is a common pathology in ALS/FTD, we asked whether the elevated *C9ORF72* aberrant splicing may be in part due to the loss of TDP-43 function. We quantified *C9ORF72* splice junctions in previously published RNA-seq data from C9 NRE^+^ neuronal nuclei sorted based on their TDP-43 abundance^39^ and did not observe a significant change in the cryptic splice site usage between TDP-43^+^ and TDP-43^−^ nuclei (**Supplementary Fig. 5**), suggesting that *C9ORF72* aberrant splicing occurred upstream or independent of TDP-43 dysfunction.

### Repeat length-dependent activation of cryptic splice sites by C9 NRE

Elevated *C9ORF72* aberrant splicing could be due to either NRE inhibiting the canonical, upstream 5ʹ splice sites or NRE activating the downstream cryptic splice sites. To distinguish between these two possibilities, we first tested whether NRE can inhibit canonical splice sites by inserting varying lengths of (GGGGCC)_n_ repeats into an intron-containing reporter mRNA^40^ (**Fig. 5a**) and quantifying the splicing efficiency by RT-qPCR. NRE-containing constructs did not show significantly decreased splicing efficiency compared to the NRE^−^ control (**Fig. 5a**), suggesting that the inhibitory effect of (GGGGCC)_n_ repeats on canonical splice sites was modest if any.

**Figure 5.**
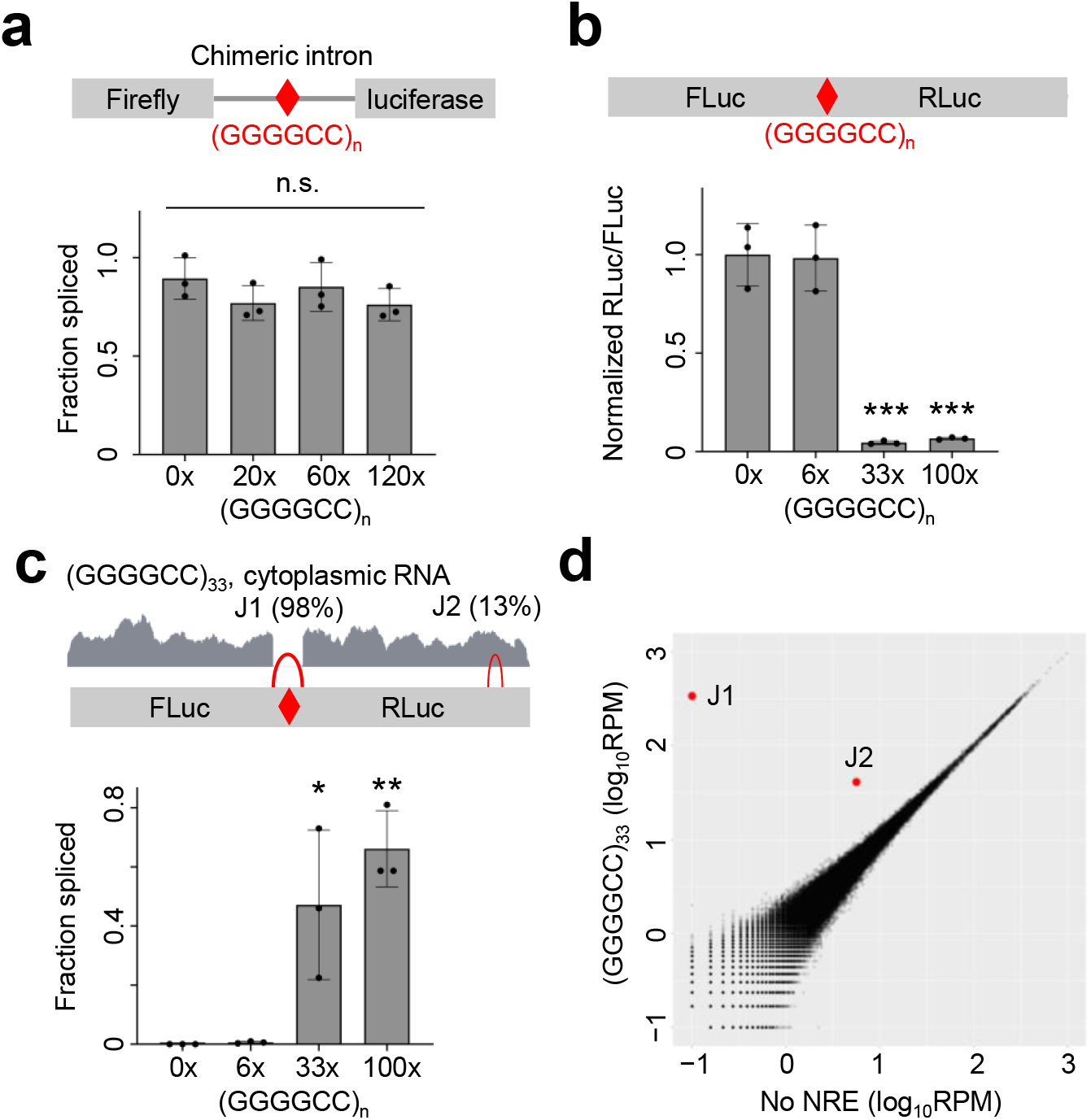
Repeat length-dependent activation of cryptic splice sites by C9 NRE. (**a**) RT-qPCR quantification of the splicing efficiency of reporter mRNAs with varying (GGGGCC)_n_ inserts within a chimeric intron. (**b**) Normalized activities of dual-luciferase with varying (GGGGCC)_n_ inserts between FLuc and RLuc coding sequences. ***, p<0.001, two-tailed t test. (**c**) *Top*, RNA-seq read coverage and splice junctions of the reporter containing (GGGGCC)_33_. Splicing efficiencies of the J1 and J2 splice junctions are shown. *Bottom*, RT-qPCR quantification of the splicing efficiency of the J1 splice junction in reporters with varying (GGGGCC)_n_ inserts. *, p<0.05; **, p<0.01, two-tailed t test. (**d**) RNA-seq quantification of all splice junctions, comparing cytoplasmic RNAs between HEK293T cells expressing reporters with no NRE and (GGGGCC)_33_. J1 and J2 splice junctions are shown in red.

To test whether C9 NRE is sufficient to activate cryptic splice sites, we inserted varying lengths of (GGGGCC)_n_ repeats into an intron-less dual luciferase reporter construct (**Fig. 5b**). Without cryptic splicing, these reporter mRNAs would be translated into firefly luciferase (FLuc) and Renilla luciferase (RLuc) at equal amount. Any cryptic splicing would potentially disrupt the coding sequence of either or both luciferases. Compared to the NRE^−^ control, (GGGGCC)_6_ (normal length) did not significantly alter the RLuc/FLuc ratio. However, (GGGGCC)_33/100_ repeats above the pathological threshold drastically decreased the RLuc/FLuc ratio (**Fig. 5b**). In contrast, the same (GGGGCC)_33_ reporter, when in vitro transcribed and transfected as mRNAs, did not significantly change RLuc/FLuc ratio compared to NRE^−^ control (**Supplementary Fig. 6**), consistent with the possibility of the reporter being aberrantly splice when transcribed in cells.

We proceeded to identify the repeat-activated cryptic splicing events by using RNA-seq. In the (GGGGCC)_33_ reporter, we detected two aberrant splice junctions, J1 and J2, both of which disrupted the Renilla luciferase coding sequence (**Fig. 5c**). Remarkably, nearly all (98%) reporter mRNAs contained the J1 junction (**Fig. 5c**, top), which explained the drastic decrease in RLuc activity. We performed RT-qPCR to quantify the J1 splicing efficiency comparing between reporter constructs with varying lengths of (GGGGCC)_n_ repeats. Consistent with our dual-luciferase assay and RNA-seq results, the disruptive J1 splicing was largely undetectable in NRE^−^ and (GGGGCC)_6_ reporters but substantially elevated when (GGGGCC)_n_ repeats exceeded the pathological threshold (33x and 100x) (**Fig. 5c**, bottom). C9 NRE has previously been associated with global defects in mRNA splicing, in part due to the sequestration of a variety of splicing factors by the C9 NRE repeat RNAs^27,41,42^. To test whether the aberrant splicing of NRE-containing reporters may be an indirect consequence due to global splicing changes, we quantified the abundance of all splice junctions within endogenous mRNAs. Only J1 and J2 were substantially increased when comparing between cells expressing the NRE^−^ and (GGGGCC)_33_ reporters, while nearly all endogenous splice junctions remained unchanged (**Fig. 5d**). These results indicated that (GGGGCC)n NRE activation of cryptic splice sites occurred predominantly in cis.

## DISCUSSION

A surprising feature of NREs associated with dominantly inherited diseases is their apparent promiscuity in producing repetitive and often more than one toxic polypeptide. The mechanism by which *C9ORF72* intronic NREs can access the cytoplasmic translation machinery has remained unclear. Our unbiased NRE-RAP-seq analysis in patient-derived fibroblast showed that *C9ORF72* intronic NRE was predominantly retained as part of an extended exon 1 in an aberrantly spliced mRNA. We found that the three most abundant splice junctions in C9 NRE-containing transcripts all used the same canonical 3ʹ splice site of intron 1. But they differed in their 5ʹ splice sites. The Ex1b-Ex2 junction has previously been known as part of the main C9ORF72 mRNA variant V2 that initiates downstream of and therefore does not contain the NRE^34^. However, our results showed that a substantial fraction of C9 NRE-containing transcripts used the Ex1b splice donor, which may be in part due to its proximity to the NRE compared to the other downstream splice donors. Notably, some downstream cryptic 5ʹ splice sites, including those of exon 1c and 1d, have previously been observed in C9 NRE^+^ iPSCs^43^, although it was not known whether C9 NRE was part of these transcripts.

This “exonization” mechanism retains C9 NRE within a capped, polyadenylated, and spliced mRNA. These mRNA features presumably enhance its stability, nucleocytoplasmic export, and translation^38^, in line with a previous study showing that C9ORF72 RAN translation is cap- and eIF4A-dependent^44^. Consistent with our finding that most C9 NRE-containing transcript used the canonical 3ʹ splice site of intron 1, siRNAs targeting exon 2 reduced the abundance of both polyGA and polyGP in fibroblasts. Similar results were recently reported by a study using an artificial microRNA to target exon 2 in *C9ORF72* transgenic mice^37^. Notably, the same study has also shown that exon 2-targeting microRNA could reduce *C9ORF72* intron RNA abundance in the cytoplasm^37^, which is consistent with our model. While these results do not rule out the possibility that NRE-containing transcripts that fully retain intron 1 or excised intron 1 lariats might also contribute to DPR production, the abundance of these RNA isoforms, as assessed by the NRE-RAP-seq read coverage in the 3ʹ half of *C9ORF72* intron 1, appears to be exceedingly low compared to C9 NRE-exonized isoforms. Furthermore, we did not find evidence of the accumulation of C9 NRE-containing unspliced pre-mRNAs nor excised intron lariats in the nucleus, suggesting that the prevalence of C9 NRE-exonized isoforms in the cytoplasm is not merely due their higher efficiency in nucleocytoplasmic export. Rather, exonization is the predominant biogenesis pathway for C9 NRE-containing transcripts.

Widespread aberrations in pre-mRNA processing have been shown as a pathological hallmark of C9 ALS/FTD, with thousands of mRNAs differentially spliced between C9 NRE^+^ and NRE^−^ ALS/FTD brains^27,42^. Previous studies have suggested that transcriptome-wide changes in splicing may be a consequence of one or more splicing factors being sequestered by either NRE RNA foci or insoluble protein aggregates^27,41^. We and others have shown that global deficits in translation-dependent mRNA surveillance caused by arginine-rich DPRs also contribute the accumulation of mis-spliced mRNAs^21,30^. In addition to mis-splicing being a downstream pathology, our results indicate that aberrant mRNA splicing also underlies the biogenesis of C9 NRE-containing RNA. Cryptic splice site activation by C9 NRE above the pathological threshold may be partly due to its recruitment of one or more splicing regulators known to interact with (GGGGCC)_n_ sequence such as hnRNP F/H^27^ and SRSF1^ref.45^. Alternatively, ectopically expressed (GGGGCC)_n_ NRE RNAs have previously been shown to be enriched near paraspeckles^46^, which are also concentrated in splicing factors. The exact contributions of these and other possible factors in C9 NRE exonization thus represent an important question for future investigation.

Our finding of a substantial fraction of NRE-containing reporter mRNAs being aberrantly spliced also echoed a recent study showing similar results using (CAG)_n_ RAN translation reporters^47^. Together, these findings strongly caution the use of NRE expression constructs such as RAN translation reporters and the interpretation of results when using these constructs. To avoid mis-interpretation due to cryptic promoter- and/or aberrant splicing-associated artifacts, thorough validation of these constructs such as RNA sequencing should be conducted to ensure the correct processing of NRE RNAs and the absence of cryptic splicing products.

As mentioned, several other NRE mutations in their native genomic contexts have been associated with aberrant splicing of their host pre-mRNA, leading to either gain- or loss-of-function consequences. (CAG)_n_ NRE in huntingtin (*HTT*) exon 1 caused aberrant splicing and pre-mature cleavage and polyadenylation in both knock-in mouse models of HD and patient tissues^48^, resulting in truncated yet still translatable mRNAs for polyQ synthesis. In addition to transcriptional silencing, mis-splicing of FMR1 mRNAs in individuals with fragile X syndrome further reduces FMRP abundance^49^. Furthermore, multiple intronic NREs have been shown to suppress canonical splicing and to cause the retention of their respective host introns^50,51^, including (CCTG)_n_ in *CNBP* intron 1 (DM2), (CTG)_n_ in *TCF4* intron 3 (Fuchs endothelial corneal dystrophy), and (GGCCTG)_n_. in *NOP56* intron 1 (SCA36). While our findings on C9 NRE suggested a distinct exonization mechanism, repeat-induced aberrant splicing may represent a common feature of NREs and a critical determinant of their pathogenicity.

## METHODS

### Cell culture

Fibroblasts and iPSCs were obtained from the National Institute of Neurological Disorders and Stroke (NINDS) Human Cell and Data Repository. After thawing, fibroblasts were cultured in Dulbecco’s Modified Eagle Medium (DMEM) supplemented with 15% heat-inactivated fetal bovine serum (FBS) and 1x non-essential amino acids (NEAA) (ThermoFisher) and were typically passaged at a 1:6 dilution ratio.

iPSCs were grown in six-well plates coated with matrigel (Corning, 354277) in mTeSR Plus media (Stem Cell Technologies, 100-0276). Upon reaching 90% confluence, iPSCs were either differentiated or passaged using Accutase (Sigma-Aldrich, A6964) and resuspended in media containing Y27632 (10 µM, Tocris, 1254).

### Motor neuron differentiation

iPSC differentiation into motor neurons was performed as previously described^52^. Upon reaching 90% confluency (day 0), iPSCs were dissociated using Accutase (Sigma-Aldrich, A6964) for 5 min at 37°C. The cells were then centrifuged at 200 rcf for 5 min and resuspended at a density of 2×10^5^ cells/ml in ultra-low attachment 6-well plates (Sigma, CLS3471) using N2B27 medium (50% Advanced DMEM/F12 (ThermoFisher, 12634010), 50% Neurobasal Medium (ThermoFisher, 21103049), supplemented with 1% Pen/Strep (ThermoFisher, 15140122), Glutamax (ThermoFisher, 35050061), and 0.1mM 2-mercaptoethanol (ThermoFisher, 31350010), B27 (ThermoFisher, 17504044), and N2 (ThermoFisher, 17502048)) supplemented with Y27632 (10 µM, Tocris, 1254), FGF2 (10 ng/ml; ThermoFisher, 13256029), SB431542 (20 µM; Tocris, 1614/10), LDN193189 (0.1 µM; Tocris, 6053/10), CHIR 99021 (3 µM; Tocris, 4423/10), and ascorbic acid (10 µM, Tocris, 4055/50). At days 2 and 4, embryoid bodies were visible, and cells were centrifuged at 200 rcf for 5 min and resuspended in N2B27 with the same supplements as on day 0 except for the removal of Y27632 and addition of all-trans retinoic acid (100 nM; Tocris, 0695/50) and SAG (500 nM; EMD Calbiochem, 566660). At day 7, the embryoid bodies were centrifuged at 200 rcf for 5 min and resuspended in N2B27 supplemented with BDNF (10 µg/ml; PeproTech, 450-02), ascorbic acid, all-trans retinoic acid, and SAG. Media were changed on days 9 and 11 with N2B27 with the same supplements as day 7 but with the addition of DAPT (10 mM; Tocris, 2634/10). At day 14, the cells were resuspended in the same media as day 9, but with the addition of GDNF (10 µg/ml; PeproTech, 450-10).

On day 16, the embryoid bodies were dissociated in 3 mL of pre-warmed 0.05% Trypsin-EDTA (ThermoFisher, 25300062) and DNase I (Fisher, NC9199796) for 1 hour at 37 °C. After adding an equal volume of FBS (ATCC, SCRR-30-2020) and DNase I, the cells were centrifuged at 200 rcf for 5 min. After collecting the supernatant, 1 ml cold PBS (Thermo Fisher, 20012027) was added to the pellet, which was then triturated gently 15 times and let to rest for 5 min. The supernatant was collected, and this process was repeated three times with 2 mL of PBS. The pooled collected supernatant was filtered through a 40 µm cell strainer (Fisher, 08-771-1) into a tube containing 2 ml cold PBS. The cells were centrifuged at 200 rcf for 5 min, resuspended in 1 ml cold PBS, and counted. iPS-MNs were resuspended in NDM medium (Neurobasal Medium supplemented with Glutamax, 0.1mM 2-mercaptoethanol, N2, Pen/Strep, MEM NEAA, ThermoFisher, 11140050), supplemented with GDNF (10 µg/ml), CNTF (10 µg/ml; PeproTech, 450-13), BDNF (10 µg/ml), IGF-1 (10 µg/ml, PeproTech, 100-11), retinoic acid (100 nM), ascorbic Acid (10 µM), and plated in six-well plates pre-coated with PDL (Corning, 354413) coated with laminin (10 µg/mL; Invitrogen, 23017-015) at a density of 2 x 10^6^ cells/ml. NDM media with supplements was replaced every 2-3 days.

### NRE-RAP-seq

5ʹ biotinylated (CCCCGG)_3_ ASO was synthesized by Integrated DNA Technologies. NRE-RAP was performed similarly as the previously described RAP method^36^ with modifications. 15 pmol biotin-(CCCCGG)_3_ ASO was first denatured at 85 °C for 3 min and immediately transferred on ice, before being mixed with 2 µg input RNA in 300 µl preheated LiCl Hybridization Buffer (10 mM Tris-HCl pH 7.5, 1 mM EDTA, 500 mM LiCl, 1% Triton X-100, 0.2% SDS, 0.1% sodium deoxycholate, and 4 M urea) and incubated at 55 °C for 2 hours on a thermomixer. 200 µl MyOne Streptavidin C1 Dynabeads (ThermoFisher) were washed twice with 100 µl LiCl Hybridization Buffer before being added to the sample and incubated at 37 °C for 30 min while shaking. After incubation, the beads were magnetically separated, washed three times in 250 µl Low Stringency Wash Buffer (1× SSPE, 0.1% SDS, 1% NP-40, and 4 M urea), and once in 250 µl High Stringency Wash Buffer (0.1× SSPE, 0.1% SDS, 1% NP-40, and 4 M urea). All washing were performed at 58 °C for 3-10 min. After being washed twice in 200 µl RNase H Elution Buffer (50 mM Tris-HCl pH 7.5, 75 mM NaCl, and 3 mM MgCl_2_; just before use, add 0.125% N-lauroylsarcosine, 0.025% sodium deoxycholate, and 2.5 mM TCEP), RNAs were eluted by mixing the beads with 21µl RNase H Elution Buffer and 4 µl RNase H (New England Biolabs) and incubated at 37 °C for 30 min while shaking. After collecting the RNA eluate, the beads were eluted again with 25 µl LiCl Hybridization Buffer at 37 °C for 5 min with shaking. Two eluates were combined, cleaned up by using Monarch RNA Cleanup Kit (New England Biolabs), and finally eluted in 30 µl nuclease-free water. After ribosomal RNA depletion (Takara, 634846), RNA-seq libraries were constructed using SMARTer Pico-Input stranded RNA-seq kit (Takara, 634836). For input RNA samples, RNA-seq libraries were constructed using a KAPA RNA HyperPrep Kit with RiboErase (Roche). All libraries were sequenced on an Illumina NovaSeq 6000 according to the manufacturer’s protocols.

### PolyGA and polyGP abundance measurement

C9 NRE^+^ fibroblasts at ∼25% confluency in 10 cm dishes were transfected with either a non-targeting siRNA control or a mix of two C9ORF72 siRNAs (10nM each; Dharmacon) targeting exon 2, by using Lipofectamine RNAiMAX (ThermoFisher) according to the manufacturer’s instructions. Half the media were replaced 24 hours after transfection and once every 3-4 days thereafter. Eight to ten days after transfection, cells were trypsinized, wash once with PBS, resuspended in Lysis Buffer (50 mM Tris-HCl, pH 7.4, 300 mM NaCl, 5 mM EDTA, 1% Triton X-100, supplemented with 2% SDS and 1x phosphatase and protease inhibitors (Millipore) before use), and let sit on ice for 10 min. Lysates were sheared by using a Bioruptor (Diagenode) with 10x 30s on/30s off cycles at 4°C and centrifuged at 16,000 g for 20 min. The supernatants were collected, and their protein concentrations were measured by using BCA assays (Pierce).

PolyGP and polyGA abundance in the supernatants were measured using Meso Scale Discovery (MSD) sandwich immunoassays. In brief, to measure polyGP, an affinity purified monoclonal polyGP antibody (TALS 828.179) generated by Target ALS was used as both the capture antibody (biotinylated TALS 828.179, 2 ug/ml), and the detection antibody (sulfo-tagged TALS 828.179, 2 ug/ml)^ref.53^. To measure polyGA, an affinity purified monoclonal polyGA antibody (clone 5E9, Cat. #MABN889, Millipore Sigma) was used as both the capture antibody (5E9, 2 ug/ml), and the detection antibody (sulfo-tagged 5E9, 2 ug/ml). Samples were diluted in Tris-buffered saline to 35 μg protein per well and tested in duplicate wells. Response values corresponding to the intensity of emitted light upon electrochemical stimulation of the assay plate using the MSD QUICKPLEX SQ120 were acquired. PolyGP and polyGA abundance are normalized to internal controls.

### Subcellular fractionation and RT-qPCR

Cells grown in a 15cm dish were harvested and lysed in 260 µl RLN buffer (50 mM Tris-HCl pH 8.0, 140 mM NaCl, 1.5 mM MgCl_2_, and 0.5% (v/v) NP-40; 1000 U/ml murine RNase inhibitor (New England Biolabs) added just before use) on ice for 5 min. Cell lysates were centrifuged at 3,000 g for 5 min to pellet the nuclei. 250 µl supernatant was collected as the cytoplasmic fraction. The nuclear pellet was washed once with 250 µl RLN buffer and centrifuged at 3,000 g for 5 min. Total RNAs were extracted from each fraction by using TRIzol LS reagent (ThermoFisher) according to the manufacturer’s instructions. Residual DNA in both fractions was removed from RNA extracts by using TURBO DNA-free (ThermoFisher).

To measure the nucleocytoplasmic distribution of C9 NRE-containing transcripts, NRE-RAP was performed with equivalent amounts (i.e., from the same quantity of cells) of nuclear and cytoplasmic RNAs as input. NRE RNAs were eluted with same volume and added in the RT-qPCR reactions. RT-qPCR was performed by using Luna Universal One-Step RT-qPCR reagents (New England Biolabs), following the manufacturer’s instructions.

### Construction of NRE-containing reporters

(GGGGCC)_n_ repeats were synthesized through RepEx-PCR as previously described^54^, using 0.5 μM (each) of 5ʹ-phosphorylated (GGGGCC)_3_ and (CCCCGG)_3_ DNA oligos (Integrated DNA Technologies), Q5 Hot-Start high-fidelity DNA polymerase (New England Biolabs), and GC enhancer. RepEx-PCR products were size-selected and purified from 0.8% agarose gels. Most linear vector backbones were generated by PCR using non-phosphorylated primers. Vectors linearized by restriction enzymes were blunted and dephosphorylated to prevent self-ligation. After blunt-end ligation, ligation products were cloned in NEB Stable competent cells (New England Biolabs), which were grown at 30°C on LB plates. Individual bacterial colonies screened for the correct repeat insert size and orientation by restriction enzyme digestion and Sanger DNA sequencing, respectively.

### In vitro transcription of reporter mRNAs

T7 promoter was inserted upstream of the dual-luciferase coding sequence by using NEBuilder HiFi DNA Assembly (New England Biolabs). Verified plasmid DNAs were digested with restriction enzymes into linear double-stranded DNA templates, which were in vitro transcribed with the mRNA products being capped and polyadenylated by using HiScribe T7 ARCA mRNA Kit (New England Biolabs). mRNA products were column-purified (New England Biolabs) and verified by glyoxal agarose gels (ThermoFisher).

### Luciferase assay

HEK293T were seeded in 24-well plate at a density of 100,000 cells/well. Luciferase reporter plasmid DNAs or in vitro transcribed RNAs were transfected by using Lipofectamine 3000 (ThermoFisher) according to the manufacturers’ instructions. 24 hours after transfection, cells were washed once with PBS, and lysed in Glo Lysis Buffer (Promega) at room temperature for 5 min. 1µl lysate was diluted with 39 µl PBS before being mixed with 40 µl of Dual-Glo FLuc substrate (Promega). After 10 min incubation, FLuc activity was measured with a GloMax 20/20 luminometer (Promega). Subsequently, 40 µl of Dual-Glo Stop and Glo reagent was added to the mixture, incubated for 10 min, and measured for RLuc activity.

### NRE-RAP-seq and RNA-seq data analysis

Fastq sequences were aligned to human genome assembly GRCh38/hg38 using STAR 2.7, with the following parameters: --outFilterScoreMinOverLread 0 -- outFilterMatchNminOverLread 0 --outFilterMatchNmin 0 --outFilterMismatchNmax 2. After alignment, read deduplication was performed by using the markdup function from SAMtools. Alignment results were visualized by using Integrative Genomics Viewer. STAR-generated BAM files and the associated metadata, including brain regions and C9 NRE genotypes, were provided by the NYGC ALS consortium. For each sample, the number of reads mapping to C9ORF72 intron 1 and the numbers of reads containing each splice junction were normalized to the total number of reads mapping to the C9ORF72 gene locus (chr9:27,546,000-27,576,000). Normalized counts between C9 NRE^+^ and NRE^−^ samples were compared by using Mann-Whitney-Wilcoxon tests.

## AUTHOR CONTRIBUTIONS

S.Y. and J.U.G. conceived the project. S.Y. conducted most experiments and analyzed the results. D.W. conducted RT-qPCR analysis in iPSCs and iPS-MNs. U.S. and A.M.V. performed DPR immunoassays under the supervision of T.G., J.M.S.S. and M.A. performed iPSC culture and iPS-MN differentiation under the supervision of J.D.P., J.Z. assisted in reporter experiments. S.Y. and J.U.G. wrote the manuscript with inputs from all authors.

## ACKNOWLEDGEMENT

We thank S. Strittmatter, S. Ferguson, W. Gilbert, C. Thoreen, Z. McEachin, and members of the Guo lab for helpful discussions and comments on the manuscript. This work was supported by an NIH Director’s New Innovator Award (DP2 GM132930) and a McKnight Neurobiology of Brain Disorders Award. J. U. G. was supported by a Klingenstein−Simons Neuroscience Fellowship and is a New York Stem Cell Foundation−Robertson Investigator.

---

**Supplementary Figure 1.**
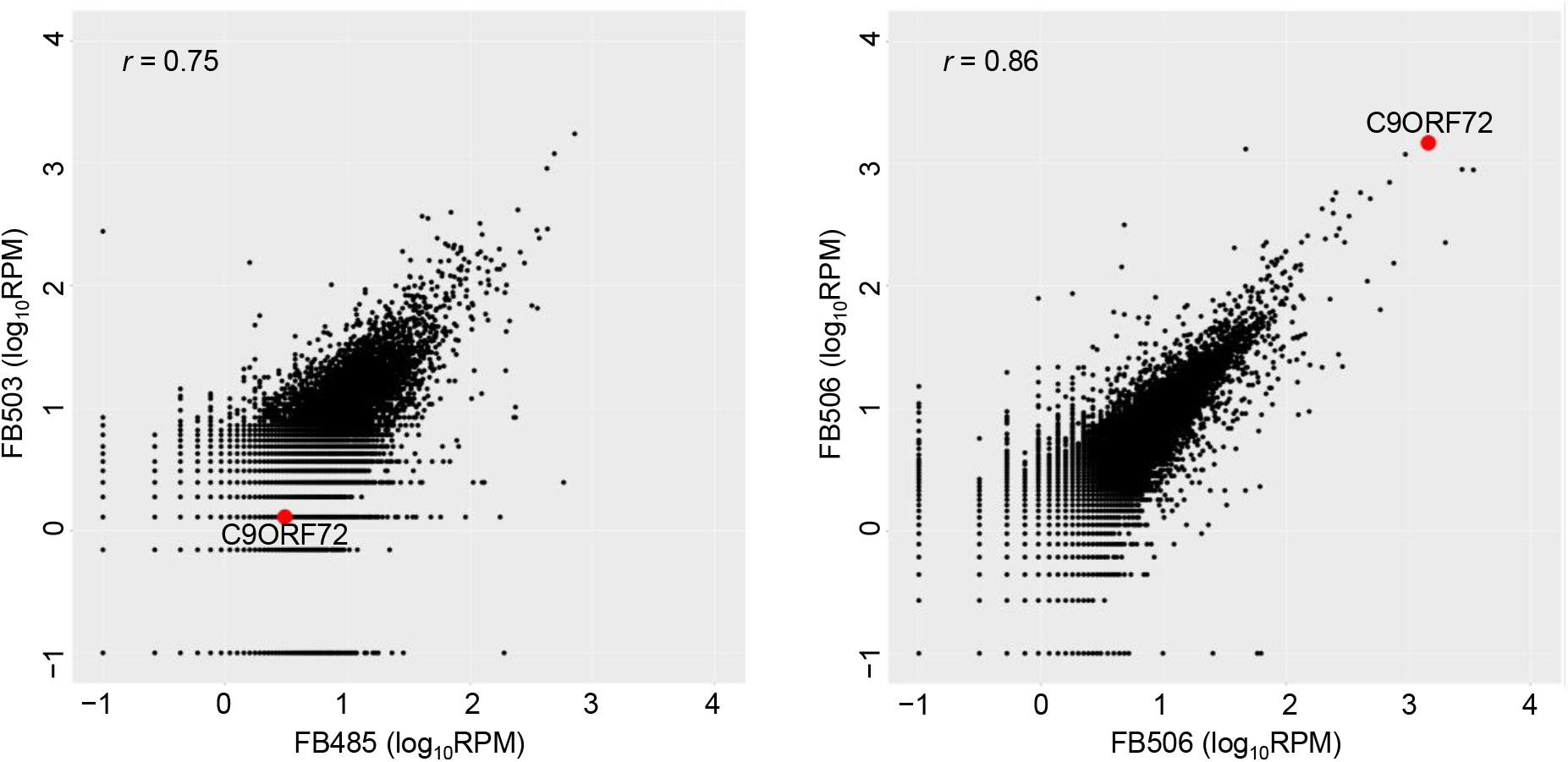
Reproducibility of cytoplasmic NRE-RAP-seq between biological replicates. Left, quantification of normalized read counts per gene, comparing two C9 NRE^−^ fibroblast cell lines. Right, quantification of normalized read counts per gene, comparing two C9 NRE^+^ fibroblast cell lines. C9ORF72 is indicated in red. Pearson’s correlation coefficients are shown.

**Supplementary Figure 2.**
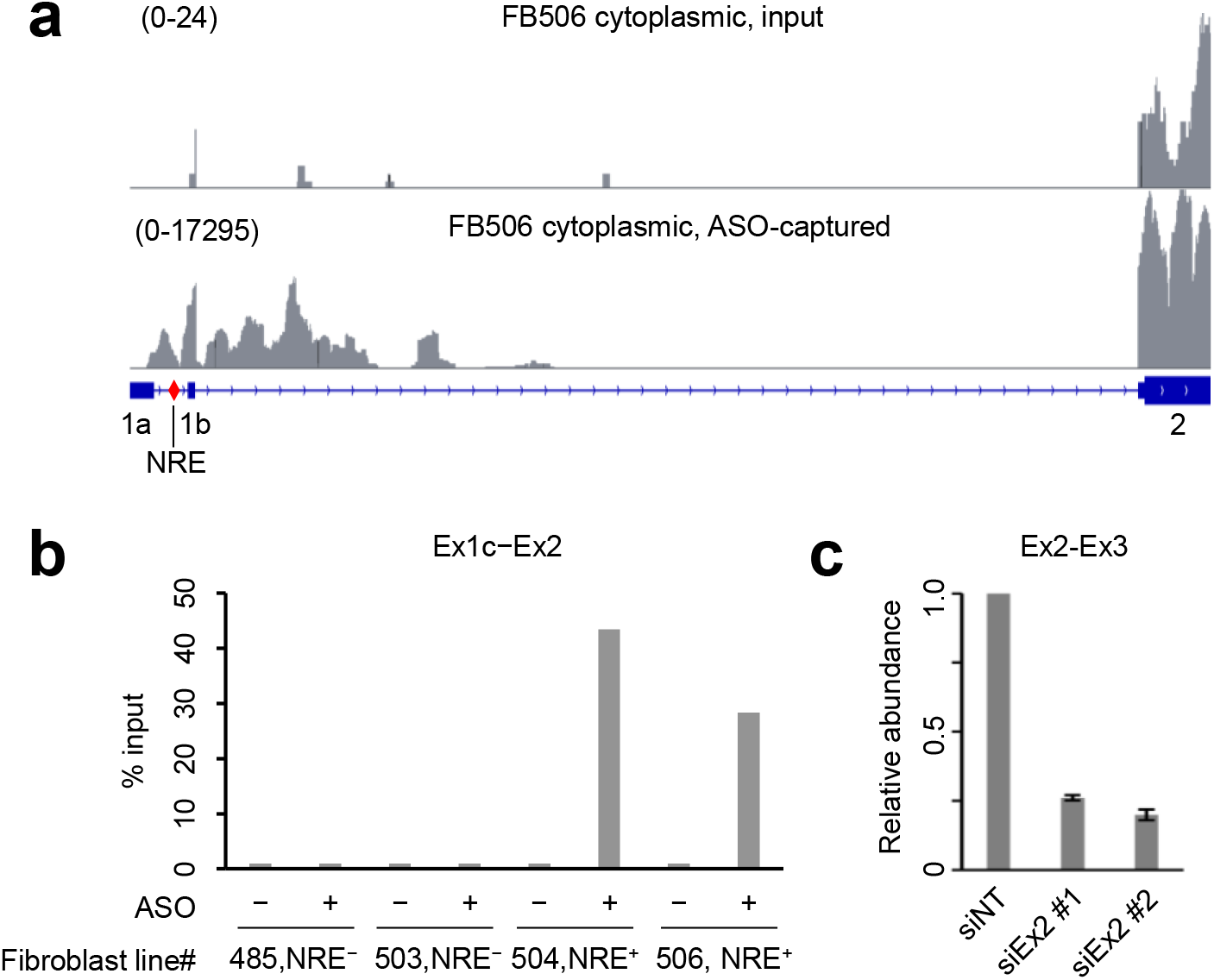
Replication and validation of cytoplasmic NRE-RAP-seq. (**a**) Read coverages of FB506 cytoplasmic RNA input (top) and ASO-captured C9 NRE-containing RNAs (bottom). (**b**) RT-qPCR quantification of the enrichment of aberrant Ex1c-Ex2 splice junction by NRE-RAP, comparing C9 NRE^+^ and NRE^−^ samples. (**c**) RT-qPCR quantification of siRNA knock-down efficiency. siNT, non-targeting siRNA. siEx2 #1 and #2, two siRNAs targeting C9ORF72 exon 2.

**Supplementary Figure 3.**
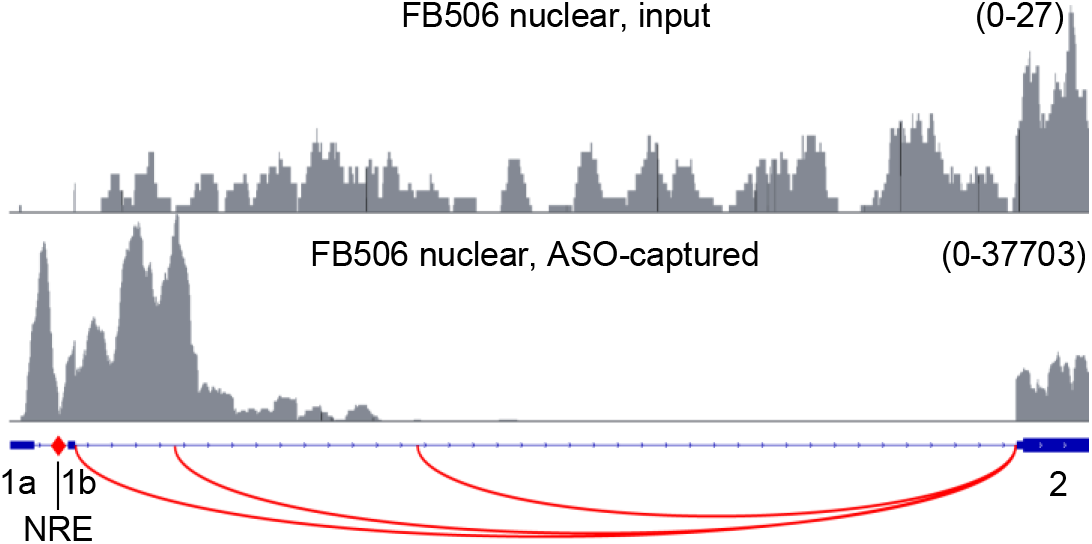
Replication of nuclear NRE-RAP-seq in FB506 fibroblasts. Read coverage of FB506 nuclear RNA input (top) and ASO-captured C9 NRE-containing RNAs.

**Supplementary Figure 4.**
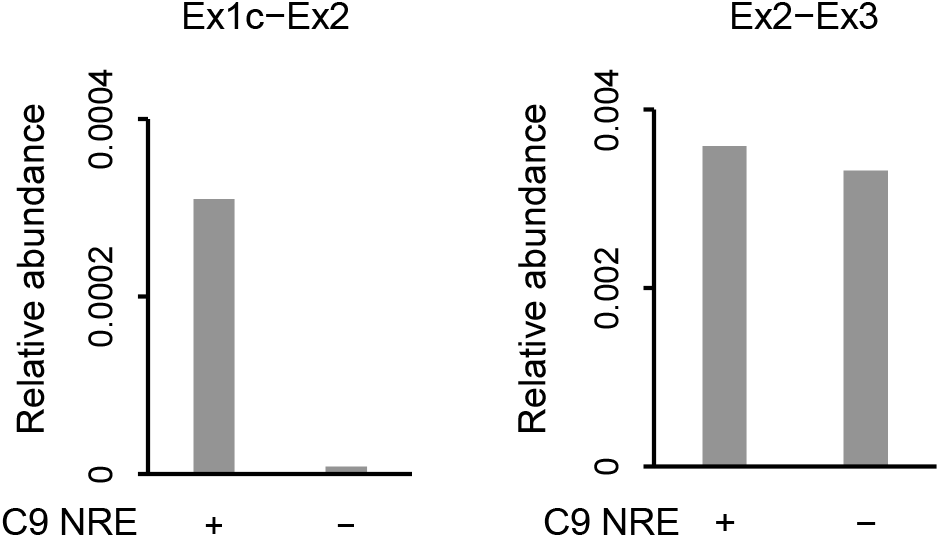
C9ORF72 aberrant splicing in C9 NRE^+^ iPSCs. RT-qPCR quantification of the aberrant Ex1c-Ex2 splice junction (left) and the canonical Ex2-Ex3 splice junction, comparing C9 NRE^+^ and C9 NRE^−^ iPSCs.

**Supplementary Figure 5.**
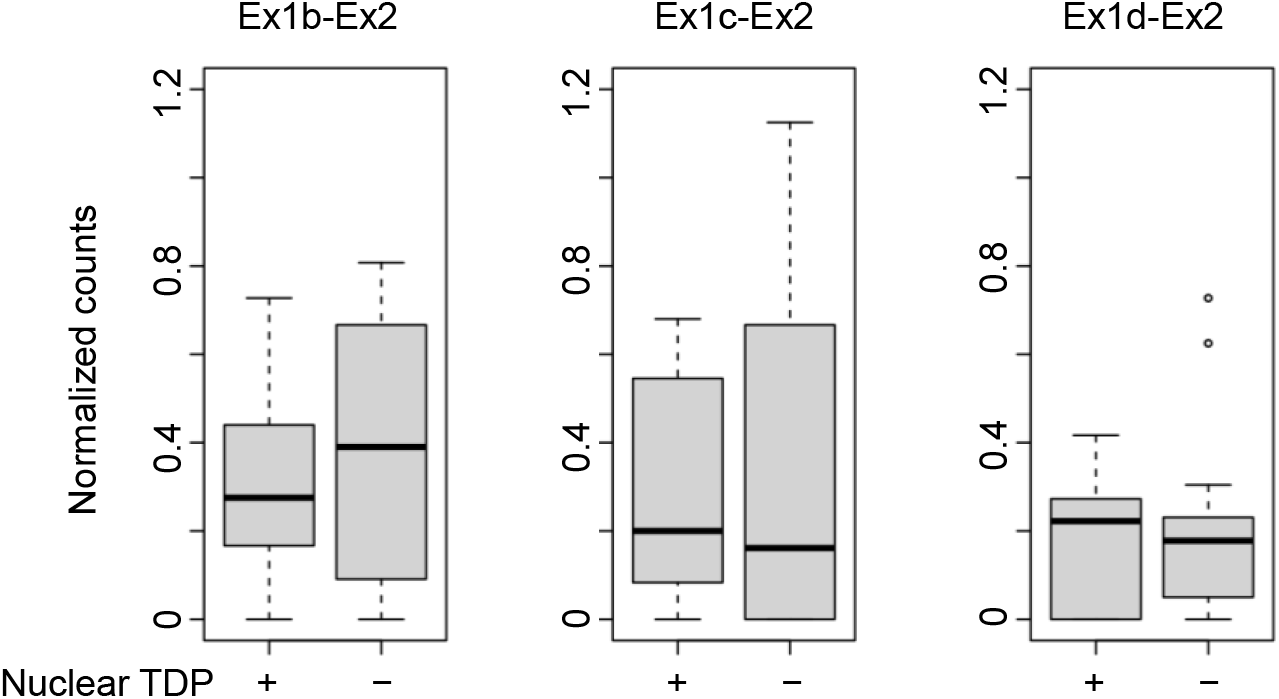
C9ORF72 aberrant splicing in C9 NRE^+^ neuronal nuclei with and without TDP-43. Normalized counts of reads spanning Ex1b-Ex2, Ex1c-Ex2, and Ex1d-Ex2 splice junctions, comparing neuronal nuclei samples with and without loss of TDP-43. RNA-seq data (GSE126543) from Liu et al. (2019) PMID: 331042469.

**Supplementary Figure 6.**
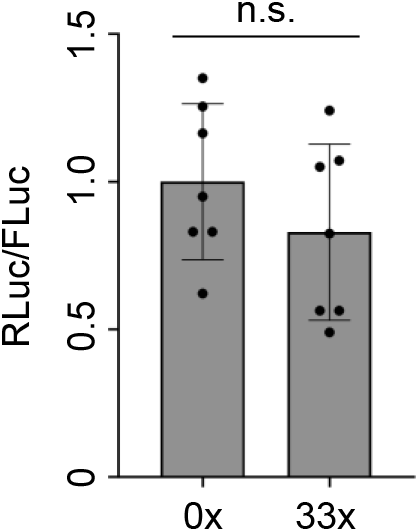
Effect of (GGGGCC)_33_ on in vitro transcribed dual-luciferase reporter mRNAs. Normalized luciferase activities from cells transfected with in vitro transcribed dual luciferase reporter mRNAs containing 0x or 33x GGGGCC repeats. n.s., not significant, two-tailed t test.

**Supplementary Table S1.**
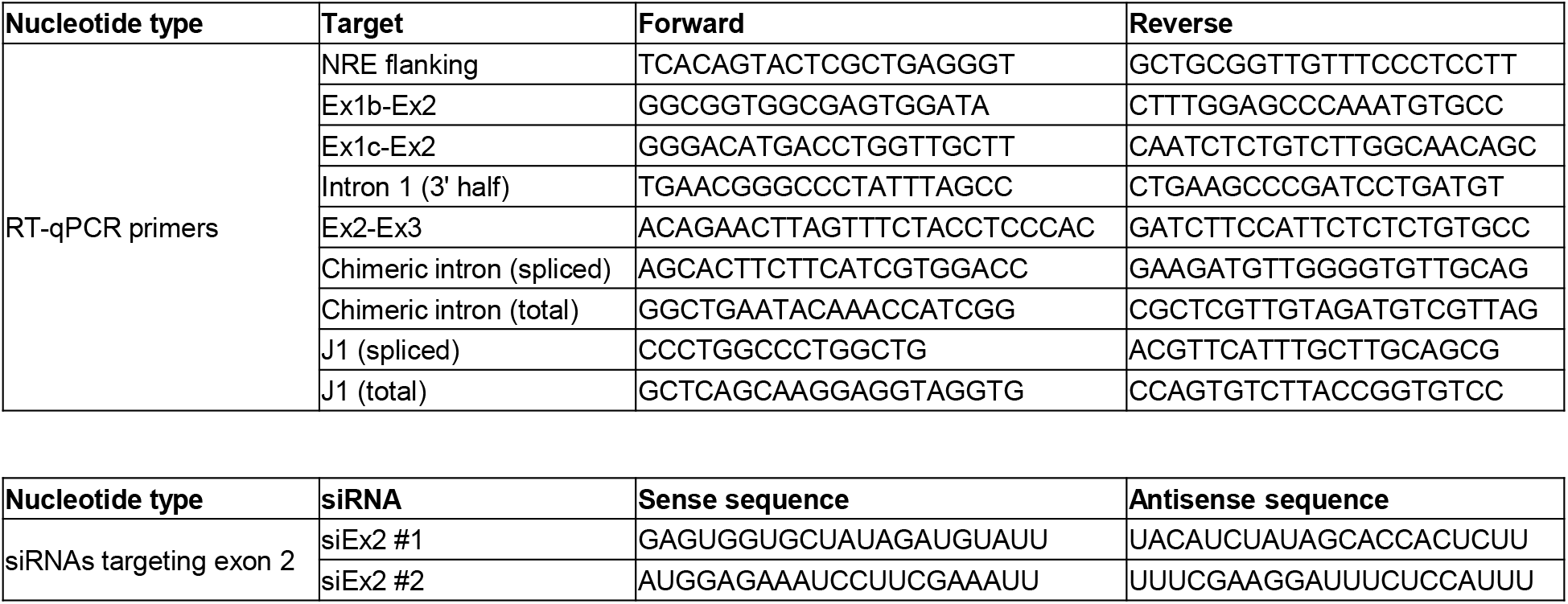
**Oligonucleotide sequences.**

